# Sheddase Targeting Chimeras (SHEDTACs) catalyze membrane target proteolysis

**DOI:** 10.64898/2026.02.06.703938

**Authors:** Brett M Garabedian

**Affiliations:** Department of Immunology and Microbiology, Scripps Research, La Jolla, CA 92037; Ablytx Inc, JLABS, 3210 Merryfield Row, San Diego, CA 92121

## Abstract

Extracellular targeted protein degradation (eTPD) has expanded access to the “undruggable” proteome but is constrained by aspects of receptor-mediated endocytosis and intracellular trafficking. This study describes **Shed**dase-**Ta**rgeting **C**himeras (**SHEDTACs**), bispecific antibodies that commandeer endogenous membrane metalloproteases (sheddases) to catalyze direct proteolysis of cell surface targets *in cis*. SHEDTACs bypass internalization requirements through enforced proximity between sheddases and substrates, enabling rapid proteolysis directly at the cell surface. Induced proximity between the immune checkpoint LAG3 and metalloprotease ADAM10 afforded nearly quantitative receptor depletion from primary human T cells. LAG3 shedding was rapid and unaffected by pharmacologic perturbation of proteasomal or lysosomal pathways, consistent with cell surface proteolysis. In a luminescent T cell reporter co-culture, SHEDTACs catalyzed LAG3 removal and enhanced T cell receptor signaling beyond conventional blocking antibodies, addressing resistance mechanisms in cancer immunotherapy. Notably, SHEDTACs depleted the non-canonical sheddase substrate, PD-1, demonstrating this approach can be expanded beyond natural protease-target pairs. SHEDTACs establish a mechanistically orthogonal eTPD platform that exploits cell surface proteolysis, offering programmable control over the cell surface proteome with broad therapeutic implications.

## Introduction

Targeted protein degradation (TPD) has expanded access to the “undruggable” proteome. Unlike modalities that require sustained target occupancy, TPD leverages transient molecular interactions to irreversibly commit therapeutic targets to endogenous degradation processes. Extracellular TPD (eTPD) strategies have emerged that include proteolysis-targeting chimeras (PROTACs)^1^, antibody-based PROTACs (AbTACs) ^2^, lysosome-targeting chimeras (LYTACs) ^3^, and a growing list of related technologies^4-17^. Modern eTPD exploits two pathways of protein turnover, (i) the ubiquitin-proteasome pathway and (ii) the endolysosomal pathway. While eTPD modalities are increasingly recognized for their ability to surmount classical barriers to drug discovery, features including tissue selectivity, degradation kinetics, and depth of depletion can be limited by receptor expression, trafficking capacity, and target-to-receptor stoichiometry. There is thus a need for additional eTPD strategies that address these limitations.

Endogenous proteolysis of membrane protein ectodomains, termed protein shedding, is a post-translational modification that governs the abundance and function of myriad membrane proteins. This process is performed by integral membrane proteases, termed protein “sheddases”, which are well-known for their broad roles in human health and disease. Among these sheddases, the ADAMs (**A D**isintegrin **a**nd **M**etalloprotease**s**) family have been characterized for their overexpression in multiple cancer types^18^. ADAM10 plays a prominent role in tumor progression through the proteolytic cleavage of protein substrates that influence cell signaling, adhesion, and migration. One substrate, the immune checkpoint Lymphocyte Activation Gene-3 (LAG3) is a potent suppressor of T cell activation that has emerged as a promising therapeutic target.

Recently termed the ‘third’ checkpoint receptor^19^, LAG3 inhibits T cell responses through binding to inhibitory ligands on T cells and antigen presenting cells (APCs), similar to the immune checkpoints Programmed Cell Death-1 (PD-1) and Cytotoxic T-Lymphocyte Antigen 4 (CTLA-4). Inhibitory LAG3 signaling is rapidly terminated through proteolytic ectodomain cleavage by ADAM10 or ADAM17 sheddases. In a therapeutic context, impaired LAG3 shedding from T cells drives resistance to anti-PD-1 therapy in murine models and clinical cohorts of melanoma and head and neck cancer (HNSCC) patients^20^. In support of these observations, simultaneous ablation of LAG3 and PD-1 was recently shown by several groups to enhance proximal T cell receptor (TCR) signaling, leading to increased proliferation and cytokine production, as well as expression of the epigenetic T cell exhaustion regulator, TOX, which is key for the persistence of exhausted T cells^21-24^. This strong synergy is evidenced by recent approval of the anti-LAG3 / PD-1 combination therapy, Opdualag^25^ and suggests a large potential for increased clinical outcomes in anti-PD-1 resistant cancers by maximizing LAG3 shedding from T cells.

In contrast to blocking strategies, induced receptor shedding offers distinct therapeutic advantages. LAG3 ligands are diverse and include major histocompatibility complex (MHC) class II molecules^26^, galectin-3^27^ (Gal-3), liver and lymph node sinusoidal endothelial cell C-type lectin (LSECtin) ^28^, fibrinogen-like protein 1 (FGL1) ^29^, α-synuclein preformed fibrils (α-syn PFF) ^30^ and, most recently, the TCR–CD3 complex^31^. Thus, blocking antibodies targeting epitopes of a single LAG3-ligand complex may not fully account for the diversity of additional, persistent LAG3-ligand complexes, including inhibitory LAG3 homodimers^32^. LAG3 shedding, on the other hand, globally depletes LAG3 from the cell surface, generating a soluble, monomeric ectodomain with relatively low affinity for its ligands^33^. Thus, strategies capable of inducing membrane protein ectodomain shedding hold immense therapeutic value as a novel eTPD strategy.

This work describes **Shed**dase-**ta**rgeting **c**himeras (**SHEDTACs**), bispecific antibodies that induce proximity between endogenous membrane proteases (sheddases) and cell surface protein targets, catalyzing membrane target proteolysis *in cis* (**Figure 1**). This concept is demonstrated through induced shedding of the immune checkpoint LAG3 by the endogenous sheddase ADAM10 on primary T cells (**Figure 2**). Using a focused SHEDTAC library, robust LAG3 shedding was achieved across diverse binding epitopes on both LAG3 and ADAM10, highlighting a degree of promiscuity that is unhindered by traditional design constraints required of earlier TPD efforts (**Figure 3**). Using a commercial T cell-based luciferase reporter, SHEDTAC treatment showed TCR signaling enhancements that outperformed anti-LAG3 blockade (**Figure 4**). Notably, the concept of induced receptor shedding was extended to the non-canonical ADAM10 substrate, PD-1 (**Figure 5**), indicating the potential to induce shedding between unnatural protease-target pairs. SHEDTACs expand the eTPD toolbox beyond proteasomal and lysosomal strategies and in this way, unlock new therapeutic opportunities to treat intractable diseases.

**Figure 1.**
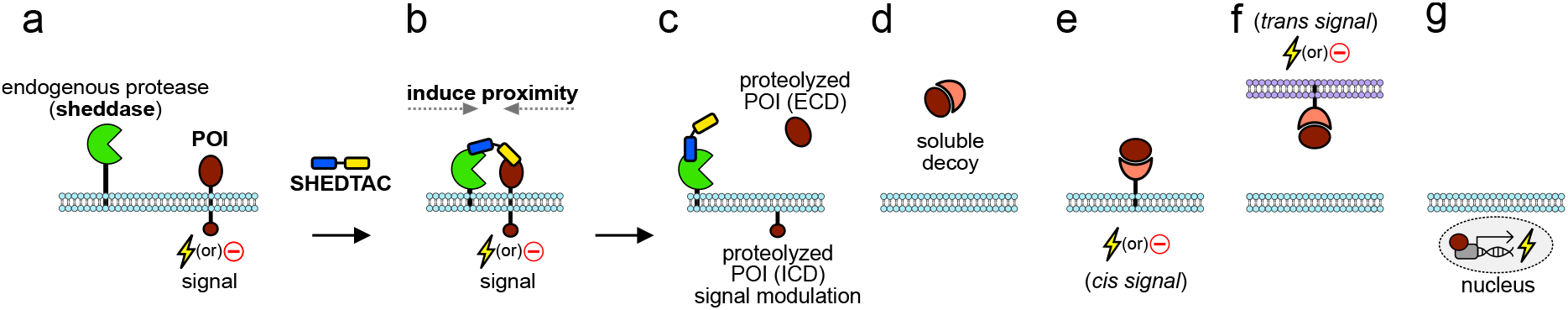
Sheddase Targeting Chimeras (SHEDTACs) catalyze membrane target proteolysis. (**a**) SHEDTACs are bispecific antibodies that induce proximity between endogenous cell surface proteases (sheddases) and a protein of interest (POI). (**b,c**) Induced proximity catalyzes POI proteolysis by the sheddase, modulating POI signaling. Proteolysis can afford fragments including extracellular domains (ECD) and intracellular domains (ICD). (**d**) Soluble ECDs act as decoys, engaging cognate ligands in solution (**d**), membrane bound in *cis* (**e**), or in *trans* (**f**) to affect downstream signaling. (**g**) ICDs are further processed into soluble species that translocate to the nucleus to affect gene transcription.

**Figure 2.**
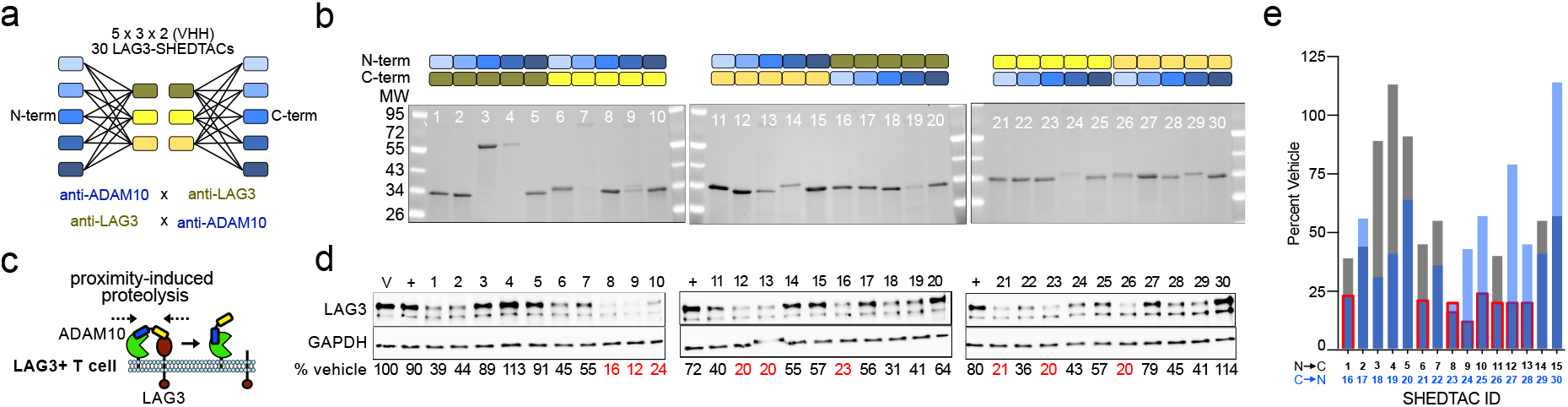
Diverse SHEDTAC configurations accelerate LAG3 shedding by ADAM10 on T cells. (**a**) Design of a SHEDTAC library spanning five anti-ADAM10 VHH and three anti-LAG3 VHH, all targeting distinct epitopes. Diverse configurations are achieved through N→C or C→N terminal VHH fusions, affording a library of thirty unique LAG3/ADAM10 bispecific combinations. (**b**) Reducing SDS-PAGE analysis of purified SHEDTACs used to treat cells in (**c,d**). (**c**) T cell surface LAG3 shedding by the protease ADAM10, which is accelerated by SHEDTACs (**d**) Western blot analysis of T cell pellets indicating levels of intact LAG3 on cells following 24h treatment, quantified in (**e**). Intensity is expressed as percent of vehicle (V). “+” indicates the addition of ionomycin (10µg/ml) to induce ADAM10 activity.

**Figure 3.**
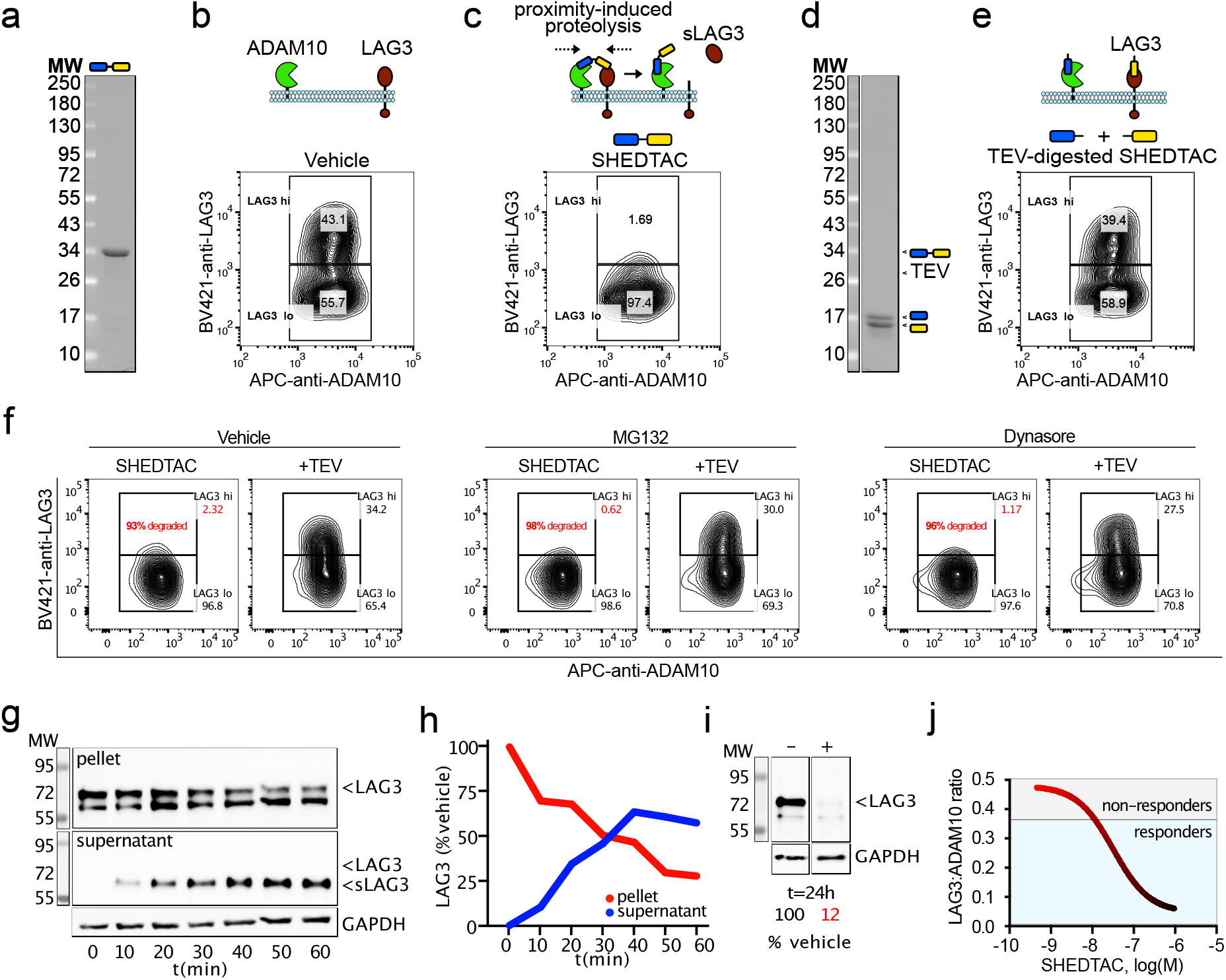
SHEDTACs drive rapid and complete LAG3 shedding by ADAM10 on T cells. (**a**) Reducing SDS-PAGE indicating SHEDTAC#8, selected for its high activity shown in **Fig2d,e**. (**b-e**) Flow cytometry contour plots showing LAG3 abundance on CD3+ADAM10+ PBMCs following 1h treatment with (**b**) vehicle, (**c**) 500nM SHEDTAC, (**d,e**) equimolar TEV-proteolyzed SHEDTAC, serving as monospecific controls. (**f**) Flow cytometry contour plots showing LAG3 abundance on CD3+ADAM10+ PBMCs treated with SHEDTAC (left) or equimolar TEV-digested SHEDTAC (right). Prior to treatment, cells were incubated for 2h at 37°C with vehicle (left plots), proteasome inhibitor (MG132, 10µM, middle plots), or lysosome inhibitor (Dynasore, 50µM, right plots). (**g**) Western blot analysis of cell pellets and conditioned cell supernatants treated with SHEDTAC, sampled every 10 minutes for 60 minutes, indicating time-dependent decreases in full-length (∼70kDa) LAG3 and concomitant increases in soluble LAG3 (sLAG3, ∼60kDa) ectodomain released into the growth medium by ADAM10. (**h**) quantification of data from (**g**) normalized to GAPDH and expressed as percent control of cell pellet at t=0. (**i**) Cells from (**g**) following 24h SHEDTAC treatment. (**j**) Ratio of LAG3:ADAM10 as determined by flow cytometry over a range of SHEDTAC concentrations.

**Figure 4.**
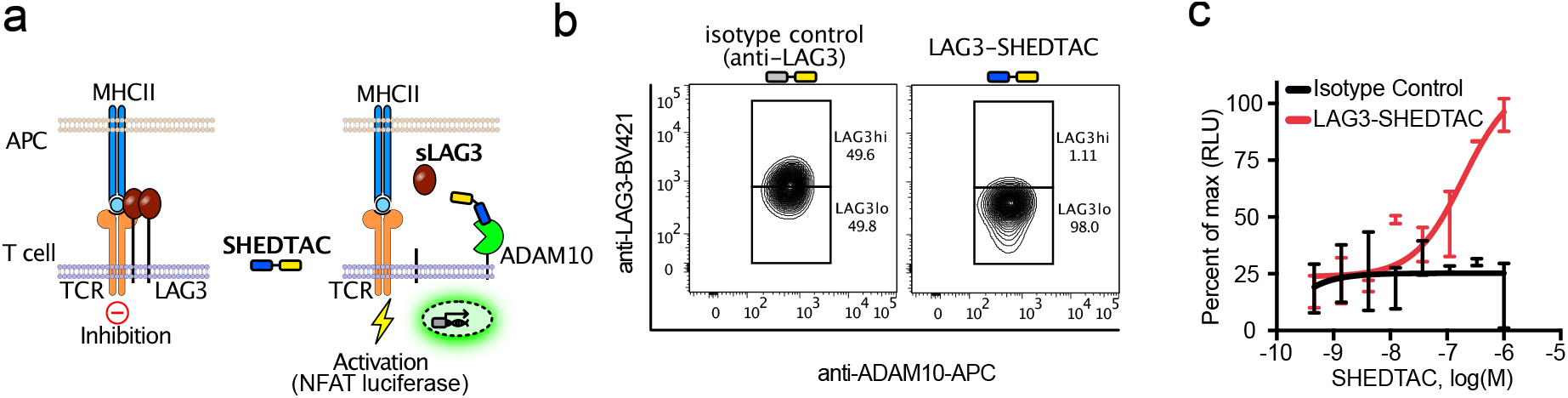
SHEDTACs enhance TCR signaling through accelerated LAG3 shedding by ADAM10. (**a**) LAG3 suppresses T cell signaling through homodimer formation, and interactions with the TCR on T cells and MHCII on antigen presenting cells (APCs) (left). LAG3-SHEDTACs catalyze LAG3 proteolysis by endogenous protease ADAM10 to restore TCR signaling and induce a luciferase reporter (right). (**b**) Flow cytometry contour plots indicating LAG3 abundance on ADAM10(+) luciferase reporter Jurkat cells treated with isotype control or LAG3-SHEDTAC. (**c**) Dose-dependent luminescence increases following treatment with SHEDTAC at the indicated concentration, illustrating enhanced TCR signaling that is afforded through LAG3 shedding. RLU = relative luminescence units

**Figure 5.**
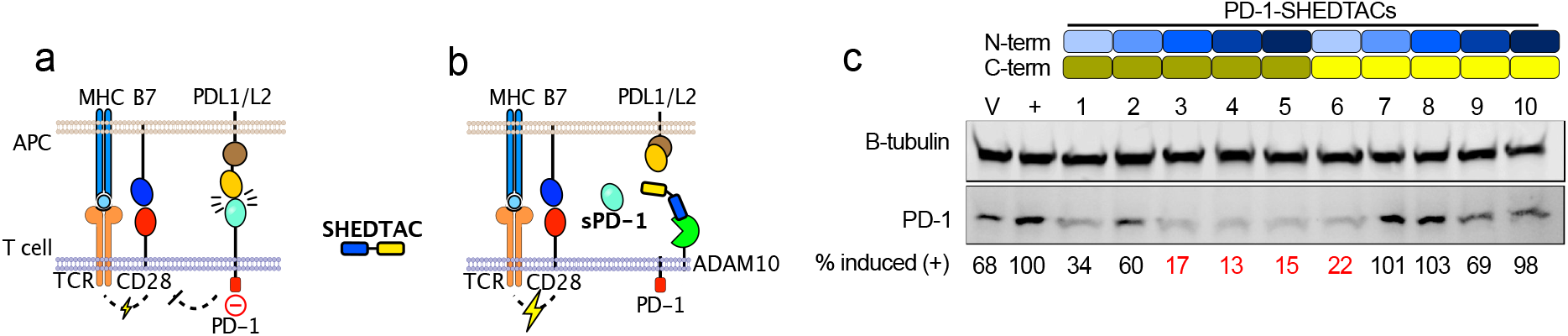
SHEDTACs catalyze proteolysis of the noncanonical sheddase substrate PD-1. (**a**) T cell activation proceeds through TCR binding to peptide-loaded MHC, followed by co-stimulator CD28 binding to its B7 ligands on APCs. The immune checkpoint PD-1 is a noncanonical sheddase substrate that broadly inhibits T cell activation. (**b**) PD-1-SHEDTACs catalyze proteolysis of the noncanonical sheddase substrate PD-1 by the integral membrane protease ADAM10 on T cells. (**c**) Western blot analysis of PD-1(+) PBMCs treated with vehicle or PMA/ionomycin to induce PD-1 expression and ADAM10/17 activity, signified by “+”. PD-1 abundance is normalized to tubulin, expressed as percent of PMA/ionomycin induced control.

## Results

Proximity-based therapeutics face unique design constraints, based on target-effector epitope choice, linker length and composition, and overall spatial geometry. In contrast to conventional therapeutic optimization, where potency tracks with affinity metrics (e.g. K_d_, K_i_), optimal proximity-based activities are realized through specific 3-D conformations that form a productive ternary complex^17^. To maximize the likelihood of productive shedding through bispecific engagement of the immune checkpoint LAG3 and the sheddase ADAM10, a focused library of non-inhibitory VHH targeting LAG3 or ADAM10 was curated (**Figure 2**). VHH spanning three unique epitopes on the LAG3 ectodomain, and five unique epitopes on the ADAM10 ectodomain, were combined in all possible orientations spanning N→C and C→N terminal VHH fusions, each separated by a glycine-serine linker region, affording a 3x5x2 matrix of thirty LAG3-SHEDTACs (**Figure 2a,b**). The resulting SHEDTACs were evaluated for their ability to accelerate LAG3 shedding by ADAM10 in the context of T cells activated from peripheral blood mononuclear cells (PBMCs, **Figure 2c**). Sheddase activity was induced on LAG3+ T cells through the addition of the calcium ionophore, ionomycin^34^. Next, LAG3+ADAM10+ cells were separately treated with each SHEDTAC from the thirty-member library and after 24h, cells were pelleted and analyzed by reducing SDS-PAGE and western blot for the presence of intact, cell surface LAG3 (**Figure 1d, Supplementary Figure S1**).

Of the thirty SHEDTACs tested, approximately twenty depleted LAG3 to below 50% of vehicle, while nine SHEDTACs achieved ≥75% shedding after 24h, as determined by diminished chemiluminescence intensity corresponding to intact LAG3 (**Figure 2d,e**). Two SHEDTACs (#3, #4) underwent errant gene duplication, affording gene products near 60kDa that did not yield productive shedding. Notably, shedding was observed across all LAG3-ADAM10 VHH binders, with activity differences deriving from N→C versus C→N configurations between matched pairs, compared in **Figure 2e**, black versus blue bars, highlighting optimal ternary complex geometries afforded through diverse SHEDTAC configurations. Subsequent experiment utilized SHEDTAC#8, chosen for its ability to drive deep LAG3 depletion.

LAG3 shedding was further evaluated using a flow cytometry assay. Following activation for 72h, CD3+ADAM10+LAG3+ PBMCs were treated with SHEDTACs for 1h at 37°C in the presence of ionomycin. Treated cells were washed, stained and analyzed by flow cytometry for the abundance of cell surface markers including CD3, ADAM10 and LAG3 (**Figure 3a-c, Supplementary Figure S2**). The two VHH domains of each SHEDTAC are separated by a glycine-serine linker region containing a tobacco etch virus (TEV) protease consensus motif. To control for artificial signal reduction that could occur through competitive epitope binding between SHEDTAC targeting domains and flow cytometry detection antibodies, SHEDTACs were TEV-proteolyzed and then applied to cells as two freely soluble constituent domains. By severing the linker region of the two targeting domains, TEV proteolysis additionally controls for the critical requirement of induced proximity between ADAM10 and LAG3 by intact SHEDTACs. Data were normalized to equimolar TEV-proteolyzed controls, and significant shedding was indicated wherever this ratio ‘x’ was x<1. In contrast, TEV-normalized shedding where x≥1 indicates low SHEDTAC activity, or VHH competition with LAG3 detection reagents, confirmed by western blot (**Supplementary Figure S3, S4**). Indeed, cells treated with TEV-proteolyzed SHEDTACs displayed cell surface LAG3 levels comparable to the vehicle control (**Figure 3d,e**), demonstrating the requirement for bispecific engagement of LAG3 and ADAM10. Each of the thirty SHEDTACs were re-analyzed by this assay, showing LAG3 depletion in good agreement with western blot analysis (**Supplementary Figure S4**).

Current eTPD strategies rely on multistep proteasomal or lysosomal trafficking to achieve protein degradation. SHEDTACs are differentiated in their mechanism of action, as they operate by a single proteolytic event constrained to the cell surface. To demonstrate this point, CD3+ADAM10+LAG3+ PMBCs were pre-treated for 2h with MG132^35^ or Dynasore^36^, inhibitors of the proteasome and lysosome, respectively. Cells were subsequently treated with SHEDTACs or equimolar, TEV-proteolyzed controls for 1h at 37°C. In all cases, robust induction of LAG3 shedding (>90%) was observed across all treatment groups, relative to TEV-digested controls, highlighting an orthogonal cell surface proteolytic mechanism that operates independent of the proteasome or lysosome (**Figure 3f**).

LAG3 shedding kinetics were evaluated using western blot and flow cytometry. Briefly, CD3+ADAM10+LAG3+ PBMCs were treated for 1h at 37°C as described above. For western blot analysis, cells were samples every 10min during a 60min time course, and supernatants and pellets analyzed separately for the presence of LAG3 ectodomain (**Figure 3g,h**). Western blot data were quantified, showing SHEDTACs accelerate LAG3 proteolysis by ADAM10 and reduce surface LAG3 levels by ∼50% after treatment for 30 minutes, and by ∼75% after treatment for 1h. Primary LAG3 sequence analysis indicates transmembrane and cytosolic domains that together comprise 8.8kDa of LAG3. It is therefore expected that a soluble LAG3 (sLAG3) ectodomain would be detected at a molecular weight that is 8.8kDa lower than intact LAG3. Indeed, cell supernatants analyzed by western blot revealed a LAG3 fragment with molecular weight that is 8.8kDa lower than full-length LAG3, consistent with the time- and dose dependent production of sLAG3 ectodomain (**Figure 3j, Supplementary Figure S5**). Cell surface LAG3 was virtually absent following 24h of treatment, as depicted in **figure 3i**.

ADAM10 abundance remained unchanged during the treatment period, and dose dependent LAG3 shedding was expressed as the mean fluorescence intensity (MFI) ratio LAG3/ADAM10 (**Figure 3j**). This LAG3/ADAM10 ratio has previously been reported as a prognostic biomarker that determines PD-1 responsiveness in HNSCC and melanoma patients. Specifically, a ratio indicating LAG3/ADAM10 > 0.38 indicates poor survival probability over a 5-year period^20^. It is therefore notable that SHEDTACs shift this ratio to LAG3/ADAM10 << 0.38, indicating the potential to phenocopy PD-1/L1 responsive patients, and potentially restore sensitivity to PD-1/L1 treatment in patients resistant to checkpoint blockade.

LAG3 suppresses T cell function through binding to MHC class II molecules, which leads to decreased TCR signaling, reduced cytokine production, and limited proliferation^37^. LAG3 shedding by ADAM10 terminates these inhibitory effects and restores T cell function^38^. To demonstrate the functional enhancements of LAG3-SHEDTAC to TCR, a commercial LAG3/MHCII blockade co-culture assay was employed. Luciferase reporter T cells were co-cultured with antigen-loaded APCs and T cell activation was determined by increased promoter-mediated luminescence (**Figure 4a**). SHEDTACs accelerated LAG3 shedding from luciferase reporter cells by ADAM10 (**Figure 4b**) to afford a dose-dependent luminescence increase, indicating enhanced TCR signaling is achieved through accelerated LAG3 shedding (**Figure 4c**). Collectively, these data demonstrated a model of LAG3 cleavage by ADAM10 that is accelerated through proximity catalyzed by SHEDTACs. Notably, LAG3 associates with the TCR on T cells *in cis*, and despite robust LAG3 shedding, no CD3 shedding was observed, nor was TCR signaling functionally diminished. These data suggest proximal, off-target shedding of LAG3-associated TCR complexes does not occur.

SHEDTACs represent a modular approach that can be reconfigured to induce proteolysis of diverse membrane targets, although it is unclear whether SHEDTACs can induce proteolysis by unnatural protease-target pairs. One such protein is the immune checkpoint receptor Programmed Cell Death-1 (PD-1) on T cells. Soluble forms of PD-1 measured in patient sera are attributed to alternative gene splicing^39, 40^. To investigate the potential to induce shedding of unnatural sheddase substrates, SHEDTACs targeting PD-1/ADAM10, termed “PD-1-SHEDTACs” were generated. These reagents were composed of two separate VHH targeting unique epitopes on the PD-1 ectodomain and five separate VHH targeting unique epitopes on ADAM10 ectodomain, separated by a glycine-serine linker region. As described for LAG3, PBMCs were stimulated with PMA/ionomycin to further induce PD-1 expression and ADAM10 activity. Each of the ten PD-1-SHEDTACs were separately applied to cells for 24h at 37°C in cell culture medium. Western blot analysis revealed increased PD-1 expression by ionomycin (“+”) that was robustly diminished by the addition PD-1-SHEDTACs spanning multiple epitopes on PD-1 and ADAM10. These data support induced proteolysis by SHEDTACs as a generalizable strategy that can be applied to non-canonical sheddase substrates.

## Discussion

Extracellular targeted protein degradation (eTPD) has expanded the druggable surfaceome by moving beyond occupancy-based pharmacology toward irreversible, event-driven target depletion. Most eTPD approaches converge on intracellular clearance, either by ubiquitin-proteasome- or endolysosomal degradation. These modalities can be practically constrained by receptor location and density, internalization kinetics, sorting capacity, and receptor/target recycling. This work describes **Shed**dase-**ta**rgeting **c**himeras (**SHEDTACs**), bispecific antibodies that induce proximity between endogenous membrane proteases and cell surface protein targets, catalyzing membrane target proteolysis *in cis*.

SHEDTACs exploit the natural process of ectodomain proteolysis, a feature of membrane proteostasis executed by endogenous metalloproteases, including the ADAMs (**Figure 1**). Unlike existing eTPD strategies that route targets for intracellular catabolism, SHEDTACs trigger cell surface proteolysis and generate defined fragments, which can be bioactive. For example, IL6R shedding produces soluble IL6R that ligates IL-6 to drive *trans* signaling through gp130^41^. In another example, MIC-A shedding releases soluble MIC-A that can dampen NKG2D on NK and CD8 T cells and promote immune evasion^42^. Endogenous shedding also acts as a potent biophysical switch. For example CD62L (L-selectin) is shed upon leukocyte activation to reduce rolling and trafficking interactions^43^. In the case of the remaining membrane stub, HER2 ectodomain shedding generates a p95HER2 membrane fragment with constitutive signaling and reduced trastuzumab engagement, driving drug resistance^44^. A classic example of downstream processing into soluble, intracellular signaling domains includes Notch or ErbB4. Once shed, these intracellular domains translocate to the nucleus and alter transcription ^45^. Protease-target pairing must therefore undergo thorough diligence to avoid undesirable, on-target effects of induced target proteolysis using SHEDTACs. For a comprehensive list of experimentally validated sheddase substrates, see https://bal.lab.nycu.edu.tw/sheddomedb/ ^46^.

This study chose LAG3-ADAM10 as an ideal target-protease pair for several reasons. LAG3 engages diverse ligands, including MHCII^26^, Gal-3^27^, LSECtin^28^, FGL1^29^, α-syn PFF^30^, TCR–CD3 complex^31^, and inhibitory LAG3 homodimers^32^. From this list, the approved anti-LAG3 antibody Relatlimab primarily targets MHCII and FGL1, resulting in tumor immune escape through alternate LAG3 ligands^47^. In contrast to checkpoint blockade, LAG3 shedding by the metalloproteases ADAM10 and ADAM17 is a biologically authentic mechanism of LAG3 antagonism that globally ablates all inhibitory ligand contirbutions^19, 23, 33, 37, 38, 47^. In a clinical context, impaired LAG3 shedding from T cells drives resistance to anti-PD-1 therapy in murine models and clinical cohorts of melanoma and HNSCC patients^20^. Thus, it was reasoned that in a PD-1 resistant context, employment of SHEDTACs to accelerate LAG3 shedding by ADAM10 could serve to restore anti-PD-1 sensitivity in patients with reduced T cell LAG3 shedding.

To this end, the current study pursued a combinatorial SHEDTAC design strategy. From a focused SHEDTAC library, nearly a third of SHEDTACs achieved ≥75% LAG3 depletion from primary T cells at 24h (**Figure 2**). Monospecific controls provide critical mechanistic confidence by showing that cleaving the bispecific into two freely soluble domains (via TEV digestion) abrogates activity, supporting an induced proximity requirement rather than epitope masking (**Figure 3a-e**). Mechanistic orthogonality to intracellular degradation is further supported by pharmacologic perturbations intended to disrupt proteasome or lysosome associated processes, which did not affect LAG3 shedding (**Figure 3f**). Kinetic data reinforce a direct shedding model, with rapid loss of full length LAG3 from cell pellets and concomitant appearance of a smaller soluble ectodomain species in supernatant (**Figure 3g-i**). In the broader eTPD context, these kinetics are notable because they suggest local enzyme-substrate turnover can drive rapid depletion without dependence on internalization and endosomal sorting, which can be rate-limiting for lysosome directed eTPD approaches^48, 49^.

LAG3 shedding was expressed as the mean fluorescence intensity (MFI) ratio LAG3/ADAM10 (**Figure 3j**). This LAG3/ADAM10 ratio has previously been reported as a prognostic biomarker that determines PD-1 responsiveness in HNSCC and melanoma patients. Specifically, a ratio indicating LAG3/ADAM10 > 0.38 indicates poor survival probability over a 5-year period^20^. It is therefore notable that SHEDTACs shift this ratio to LAG3/ADAM10 << 0.38, indicating the potential to phenocopy PD-1/L1 responsive patients, and potentially restore sensitivity to PD-1/L1 treatment in patients resistant to PD-1/L1 checkpoint blockade.

Induced LAG3 shedding afforded increased T cell activation in a luminescent reporter co-culture assay, with dose dependent enhancement of TCR signaling (**Figure 4**). This is particularly relevant for ligand-diverse checkpoint receptors such as LAG3, where biased ligand blockade may be incomplete across the spectrum of inhibitory complexes^47^. The observation that CD3 remains intact on T cells despite *cis* association with LAG3 also argues that induced sheddase recruitment does not necessarily cause proximal bystander cleavage under the tested conditions, although this will require broader proteomic profiling across cell types, reagents, proteases, and substrates.

SHEDTACs are mechanistically related to emicizumab (Hemlibra), which functions as a synthetic cofactor by bridging FIXa and FX to promote FX activation^50^. In similar way, SHEDTACs bridge endogenous sheddases with membrane protein targets to accelerate target proteolysis. A notable extension in the current work is the observation that PD-1 can be shed, and that shedding is achieved across multiple PD-1/ADAM10 epitope pairings. These data suggest induced proximity may be used to enforce proteolysis by unnatural protease-target pairs, significantly expanding the scope of eTPD, while minimizing the breadth of the “undruggable” proteome. Further adaptation of this modality to include additional protease-target pairs will generate new hypotheses and afford novel treatment strategies to address unmet patient needs.

## Methods

### Construction, expression and purification of SHEDTACs

Generalized construction and validation of SHEDTACs are shown in **Figure 2**. In brief, a total of thirty SHEDTACs that target membrane protein LAG3 and ADAM protease ADAM10 were generated. The LAG3-targeting domain and the ADAM10-targeting domain in these SHEDTAC constructs are both VHH domains. To construct these SHEDTACs, 3 VHH domains targeting LAG3 and 5 VHH domains targeting ADAM10 were used. Each VHH has a different sequence and targets a different epitope on LAG3 or ADAM10. Among the thirty SHEDTACs, 15 have the ADAM10-targeting domain at the N-terminus and the LAG3-targeting domain at the C-terminus, with the other 15 having the ADAM10-targeting domain at the C-terminus and the LAG3-targeting domain at the N-terminus. The 15 constructs in each configuration represent all possible LAG3/ADAM10 VHH pairings (3x5=15). Genes corresponding to each VHH were synthesized and clones into a pET23a vector (Novagen).

The coding sequence for each LAG3-SHEDTAC were transformed into E. coli strain BL21(DE3) (Invitrogen). E. coli harboring the pET23-LAG3-SHEDTAC plasmid were cultured overnight in sterile 15mm glass culture tubes containing 5ml of Luria Broth supplemented with 100µg/ml of carbenicillin. The following day, cells were diluted 1:100 into a sterile 500ml baffled glass flask containing 100ml of Terrific Broth supplemented with 100µg/ml carbenicillin, and grown at 37°C with shaking at 200RPM. Once cells reached an optical density (OD) of approximately 0.75, the culture was cooled to 18°C and protein expression was induced by the addition of Isopropyl ß-D-1-thiogalactopyranoside (IPTG) at a final concentration of 1mM. Following 16h expression at 18°C, 200RPM, cells are centrifuged at 5000 relative centrifugal force (RCF) and the cell pellet resuspended in 50ml of Dulbecco’s Phosphate-Buffered Saline (DPBS) containing 1mg/ml lysozyme. Cells were lysed by stirring at room temperature for 30min, cooled on ice, and further lysed using a sonicator (Fisher model FB50, amplitude 90, 3min continuous sonication). Following sonication, the 50ml cell lysate was centrifuged for 30min at 10,000 RCF, 4°C. The resulting supernatant was applied to nickel-nitrilotriacetic acid (Ni-NTA) resin and soluble LAG3-SHEDTAC (containing a C-terminal-hexahistidine affinity tag) was eluted with DPBS containing 500mM imidazole. Purified protein was buffer-exchanged into DPBS, boiled in reducing sodium dodecyl sulfate and analyzed by polyacrylamide gel electrophoresis.

### Cell Culture and Checkpoint Receptor Induction

To induce LAG3 (or PD-1) expression 6-well tissue culture plates (Falcon, 353046) were coated with 10µg of sterile anti-human CD3 antibody (Biolegend, 300438) in 2ml of Dulbecco’s Phosphate Buffered Saline (ThermoFisher, J61196.AP, PBS)) per well for 24h at 4°C. Coated plates were washed once with 2ml of PBS and peripheral blood mononuclear cells (PBMCs) were seeded into each well at a density of 1x10^6^ cells per ml in 2ml of 0.22µm sterile-filtered T complete growth medium, consisting of RPMI 1640 growth medium (gibco, 11875-093) supplemented with 10% v/v heat-inactivated fetal bovine serum (gibco, 10438-026), 55µM beta-mercaptoethanol, 1mM sodium pyruvate (gibco, 11360-070), MEM non-essential amino acids (gibco, 11140-050), 10mM HEPES buffer (gibco, 15630-080), 2mM L-glutamine (gibco, 25030-081), and 100 units/ml of penicillin/streptomycin (gibco, 15140-122). T cells were activated by the addition of 4µg sterile anti-human CD28 antibody (Biolegend, 302934) per well (2µg/ml final) and cells were cultured for 72h at 37°C in 5% CO_2_ atmosphere and 95% relative humidity.

### Flow Cytometry

Following 72h activation, cells were pooled into a sterile 50ml centrifuge tube (Genesee Scientific, 21-108) and pelleted by centrifugation at 350g for 5min at room temperature. Pelleted cells were resuspended in T complete growth medium to a density of 5x10^6^ cells per ml and seeded into a 96-well U-bottom tissue culture plate (Falcon, 353077) at 0.5x10^6^ cells per well in a 100µl final volume. Cells were treated by directly adding 10µl of a 1µM stock of SHEDTAC in PBS, or Tobacco etch virus (TEV)-proteolyzed SHEDTAC for a final SHEDTAC concentration of 100nM. Cells are incubated with SHEDTACs for 1h at 37°C in 5% CO_2_ atmosphere and 95% relative humidity without shaking. Following treatment, cells are pelleted by centrifugation at 350g for 5min at room temperature and resuspended in 200µl of sort buffer, consisting of 0.22µm sterile-filtered PBS containing 1% w/v bovine serum albumin (Prometheus, 25-529C) and 1mM ethylenediaminetetraacetic acid (EDTA, Invitrogen, AM9260G). Cells were similarly pelleted and resuspended in 200µl sort buffer for two total washes. Washed PBMCs were suspended in 50µl of sort buffer containing 1:400 dilutions of APC anti-human CD156c (ADAM10) antibody (Biolegend, 352706); Brilliant Violet 421™ anti-human CD223 (LAG-3) antibody (Biolegend, 369314); and BD Pharmingen™ FITC Mouse anti-human CD3 antibody (BD Biosciences, 555332) and incubated for 1h at 4°C. Stained cells were washed twice with sort buffer and analyzed on a ZE5 Cell Analyzer (Bio-Rad) for mean fluorescence intensities corresponding to cell surface CD3, ADAM10 and LAG3.

### Western Blot

Following 72h activation, cells were plated into 96-well U-bottom plates as described above under *Flow cytometry*. Cells were treated by directly adding 10µl of a 1µM stock of SHEDTAC serially diluted 2-fold in PBS. Cells were incubated with SHEDTACs for 24h at 37°C in 5% CO_2_ atmosphere and 95% relative humidity without shaking. Following treatment, cells were pelleted by centrifugation at 1,000g for 5min at room temperature and cell pellet and culture medium were collected and analyzed for LAG3 or soluble LAG3 (sLAG3). The entire cell pellet or 80µl of cell culture medium was boiled in reducing SDS buffer, resolved by SDS-PAGE (Bio-Rad 4568096) and transferred to a nitrocellulose membrane (Bio-Rad 1620233) in 20mM Tris, 2mM glycine and 20% methanol using an XCell II Blot Module (Thermo Fisher) powered at 35V for 120min at room temperature. The membrane was blocked at 4°C overnight in 5% w/v BSA in PBS + 0.1% v/v Tween-20 (PBST). The following day, the membrane was incubated for 1h with a 5ml volume of polyclonal goat IgG anti-human LAG3 (R&D Systems, AF2319) and rat anti-human GAPDH (Biolegend, 607902) each diluted 1:1000 in PBS containing 0.1% w/v BSA in PBST. The membrane was washed in PBST 3x15min and subsequently incubated for 1h with a 5ml volume of donkey anti-goat IgG-HRP (Santa Cruz, SC-2020) diluted 1:2000 in PBST, and mouse-anti rat IgG2a-AF647 (Southern Biotech, 3065-31) diluted 1:1000 in PBST. The membrane was washed 3x15min in PBST and stained with 3ml of SuperSignal Plus PICO West (Thermo Fisher). Chemiluminescence was immediately measured using a ChemiDoc MP Gel imaging system (Bio-Rad) at 60s exposure. Fluorescence at 647nm was similarly measured using the factory filter set and a 30s exposure.

### TCR Reporter Co-culture Bioassay

A T cell receptor (TCR) reporter co-culture assay was performed using the LAG-3/MHCII Blockade Bioassay (Promega) according to the manufacturer’s instructions. In brief, thaw-and-use MHCII APC Cells were thawed and resuspended in the kit Cell Recovery Medium containing the provided TCR Activating Antigen, then seeded into white, flat-bottom 96-well tissue culture plates and incubated overnight at 37°C in a 5% CO2 atmosphere and 95% relative humidity. The following day, SHEDTACs and indicated controls were prepared in kit Assay Buffer and serially diluted. Medium was removed from the APC plate and diluted test articles were added directly to APC wells. LAG-3 Effector Cells (Jurkat-derived reporter cells expressing human LAG-3 and a TCR-pathway–responsive luciferase reporter) were thawed, resuspended in Assay Buffer per kit instructions, and added to each well to initiate co-culture. Co-cultures were incubated at 37°C, 5% CO2 for 4h. Following incubation, Bio-Glo luciferase reagent was added per kit instructions and luminescence was measured on a plate luminometer as relative luminescence units (RLU). RLU were background-subtracted and normalized as indicated in the figure.

### Flow Cytometry of Reporter Effector Cells Following Co-culture

For analysis of cell surface marker abundance on reporter effector cells following co-culture, the non-adherent effector cell fraction was collected, pelleted by centrifugation at 350g for 5min at room temperature, and washed twice in sort buffer consisting of 0.22µm sterile-filtered PBS containing 1% w/v bovine serum albumin and 1mM EDTA. Washed cells were resuspended in 50µl of sort buffer containing fluorophore-conjugated antibodies (as indicated) and incubated for 1h at 4°C. Cells were washed twice with sort buffer and analyzed by flow cytometry to quantify mean fluorescence intensities corresponding to the indicated cell surface markers.

## Acknowledgments

I gratefully acknowledge the support of Dr. James Paulson, who encouraged my independent development of SHEDTACs and related proof of concept experiments described herein.

---

**Supplementary Figure S1.**
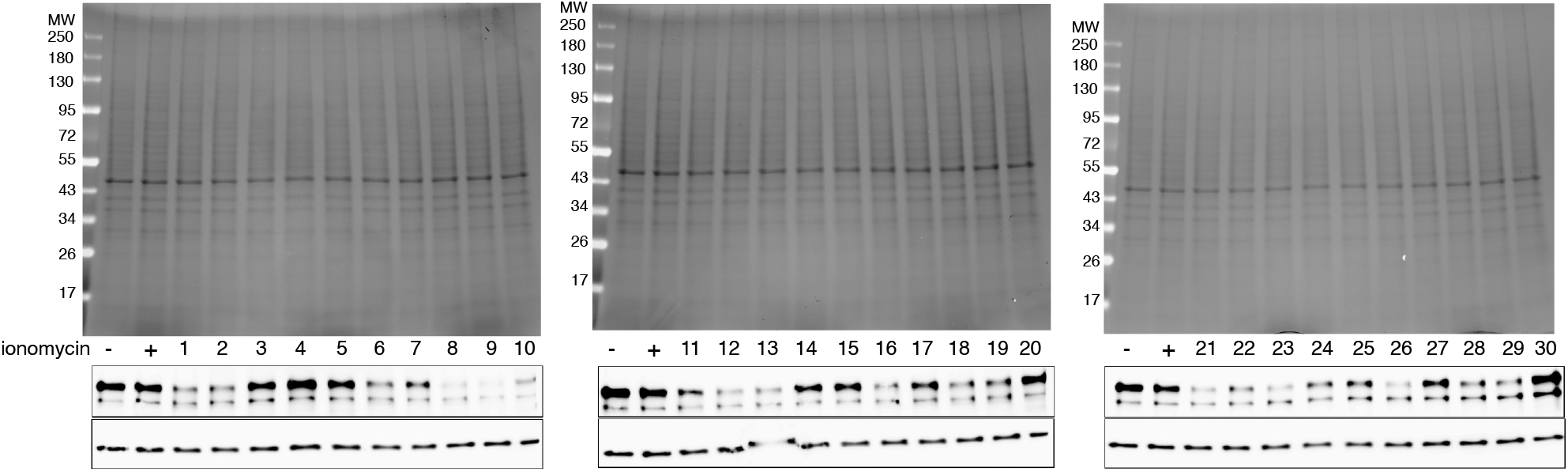
Reducing SDS-PAGE analysis of cell lysates prior to western blot analysis in **Figure 2d**

**Supplementary Figure S2.**
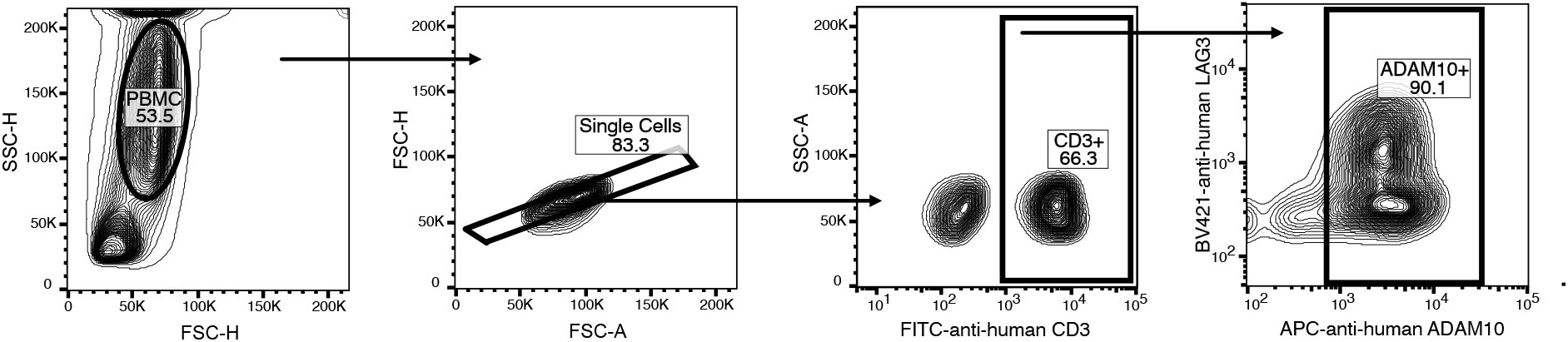
Gating strategy for activated CD3+ADAM10+LAG3+ PBMCs

**Supplementary Figure S3.**
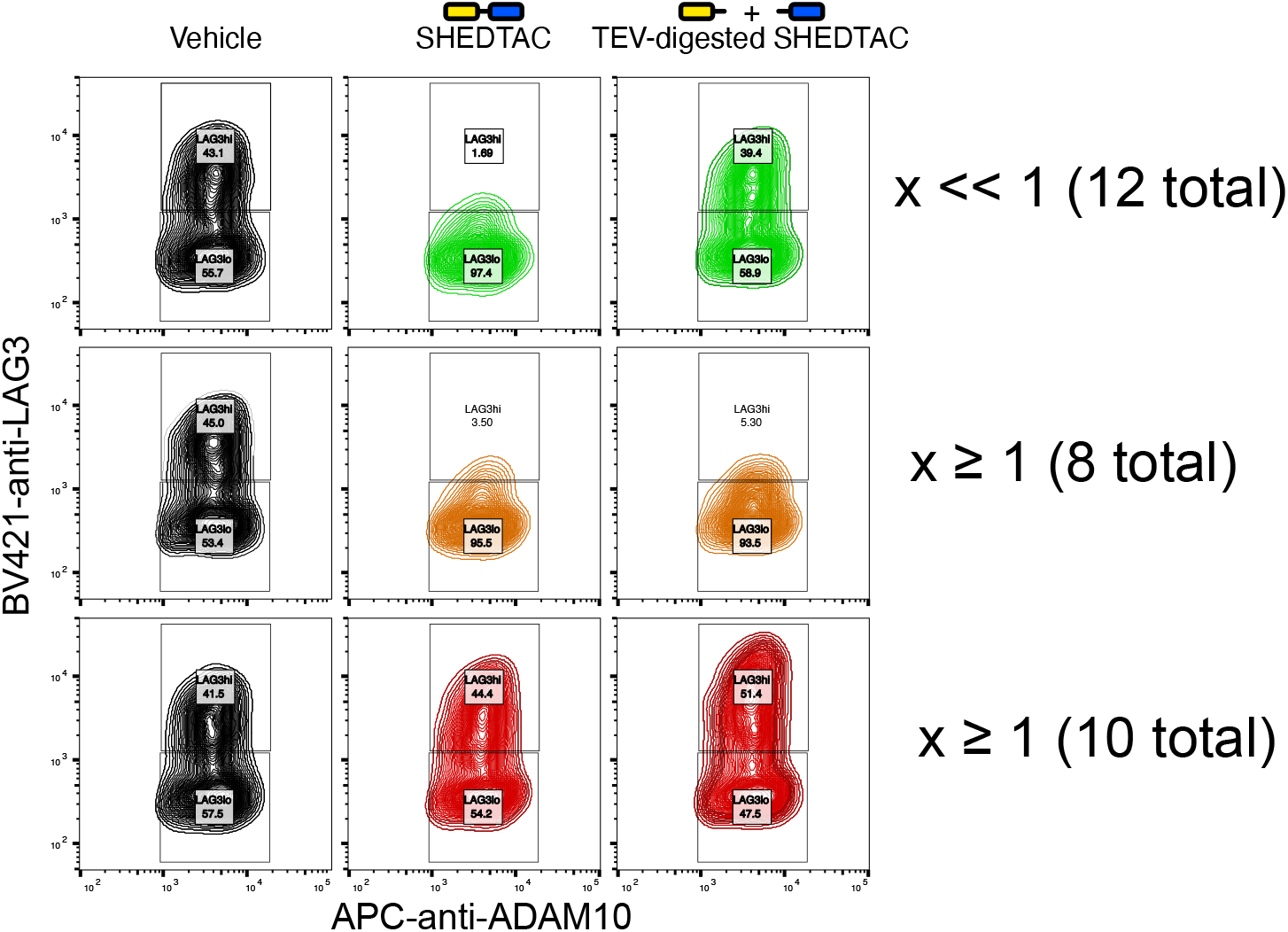
Flow cytometry TEV-normalization scheme. SHEDTACs were normalized to their equimolar TEV-proteolyzed controls, and significant shedding was indicated wherever this ratio ‘x’ was x<1. In contrast, TEV-normalized shedding where x≥1 indicates low SHEDTAC activity, or VHH competition with LAG3 detection reagents, confirmed by western blot

**Supplementary Figure S4.**
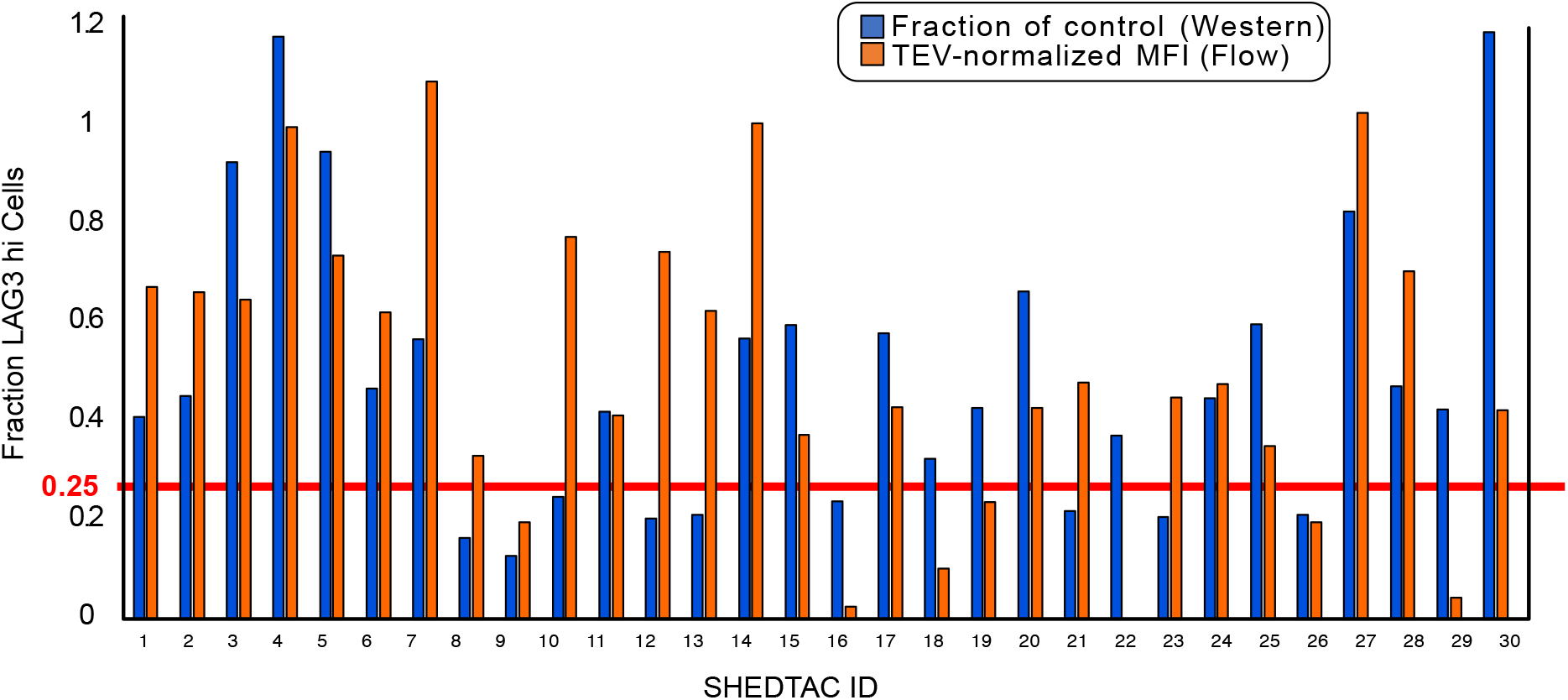
Comparison of western blot versus flow cytometry analyses to assess LAG3 shedding by ADAM10 following treatment with SHEDTACs

**Supplementary Figure S5.**
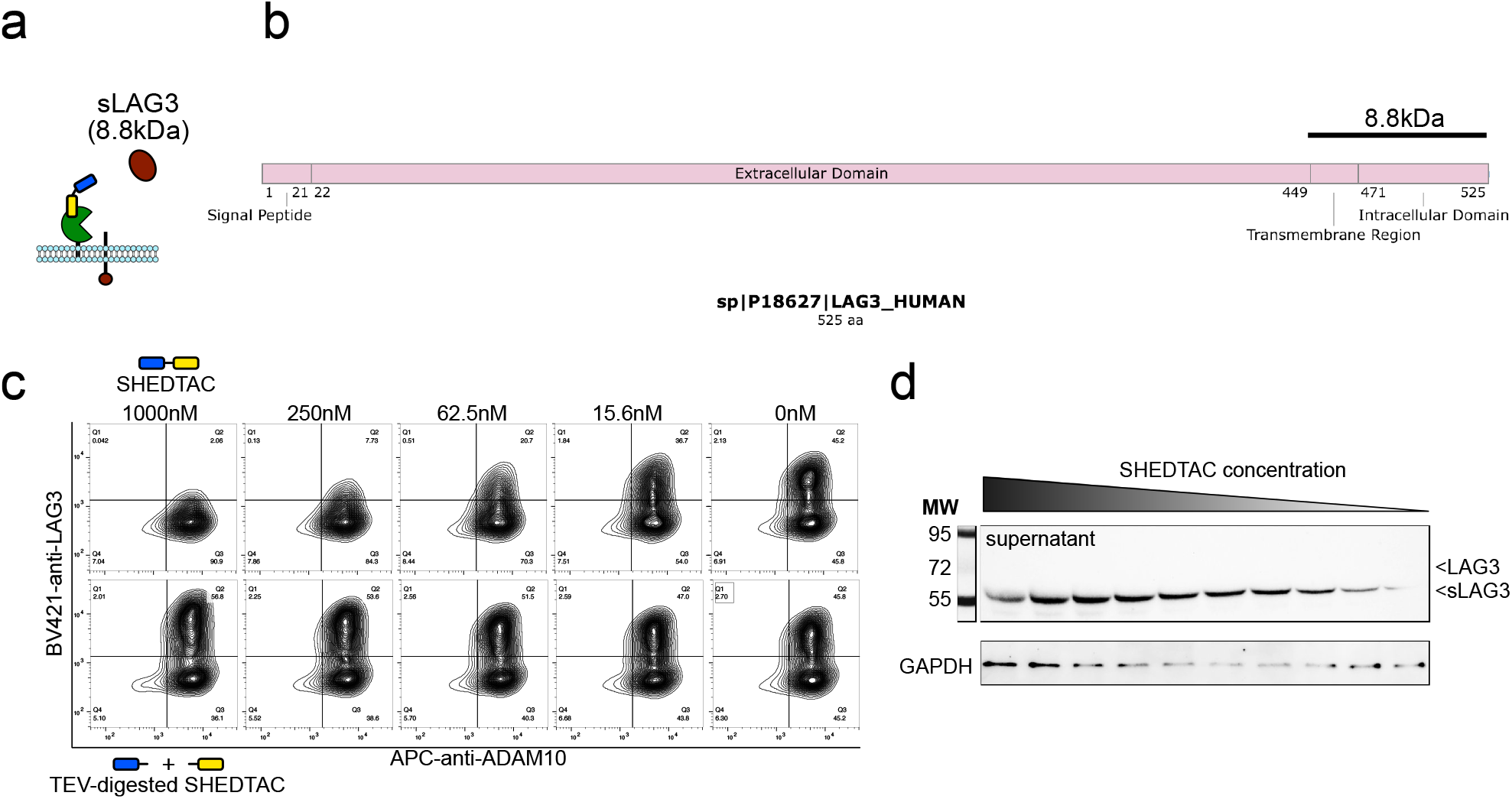
(**a**) Soluble LAG3 (sLAG3) generation through receptor shedding. (**b**) primary amino acid sequence analysis showing transmembrane and intracellular regions totaling ∼8.8kDa. (**c**) Dose-dependent loss of LAG3 abundance on T cells following treatment with SHEDTACs. Contour plots correspond to data plotted in **Figure 3j**. (**d**) Concomitant soluble LAG3 production in conditioned supernatants from cells treated in (**c**).

